# Single cell profiling in COVID-19 associated acute kidney injury reveals patterns of tubule injury and repair in human

**DOI:** 10.1101/2021.10.05.463150

**Authors:** David Legouis, Anna Rinaldi, Gregoire Arnoux, Thomas Verissimo, Jennifer Scotti-Gerber, Anna Faivre, Manuel Schibler, Andrea Rinaldi, Maarten Naesens, Kari Koppitch, Jerome Pugin, Andrew P McMahon, Solange Moll, Sophie de Seigneux, Pietro E Cippà

**Author notes:** Corresponding authors: Pietro Cippà, MD PhD, Sophie de Seigneux, MD PhD. First author equal contribution (David Legouis, Anna Rinaldi). Last author equal contribution (Sophie de Seigneux, Pietro E Cippà).

## Abstract

The cellular mechanisms of kidney tubule repair are poorly characterized in human. Here, we applied single-nucleus RNA sequencing to analyze the kidney in the first days after acute injury in 5 critically ill patients with COVID-19. We identified abnormal proximal tubule cell states associated with injury, characterized by altered functional and metabolic profiles and by pro-fibrotic properties. Tubule repair involved the plasticity of mature tubule cells in a process of cell de-differentiation and re-differentiation, which displayed substantial similarities between mouse and man. In addition, in man we identified a peculiar tubule reparative response determining the expansion of progenitor-like cells marked by PROM1 and following a differentiation program characterized by the sequential activation of the WNT, NOTCH and HIPPO signaling pathways. Taken together, our analyses reveal cell state transitions and fundamental cellular hierarchies underlying kidney injury and repair in critically ill patients.

## Main text

The development of the mammalian kidney is a finite process, ending shortly before birth in humans (and shortly after birth in mice).^1^ Thereafter, the adult kidney loses the ability to generate new nephrons, and the regenerative potential of the renal tubule depends on tissue repair. Acute kidney injury (AKI) is a common clinical syndrome associated with adverse clinical outcomes, but current standard of care continues to rely on hemodynamic optimization, avoidance of nephrotoxicity and renal replacement therapy. The application of single cell technologies generated substantial advances in the molecular understanding of kidney biology and disease,^2–7^ but the cellular mechanisms of kidney repair in critically patients are still poorly understood because kidney biopsies are rarely performed in this clinical setting and because of relevant differences among species.^8–13^

AKI emerged as a prevalent complication among patients with severe SARS-CoV2 infection: several mechanisms can contribute to the pathophysiology of COVID-19 associated AKI, but acute tubular injury secondary to hemodynamic instability, endothelial dysfunction and systemic inflammation is usually the main factor in critically ill patients.^14–16^ We characterized by single nucleus RNA sequencing (snRNAseq) kidney biopsies obtained from 5 critically ill patients with COVID-19 before planned withdrawal of resuscitation measures. All patients developed AKI in the context of severe COVID-19 with respiratory failure, cytokine storm and multi-organ involvement (**Table 1**). Notably, RT-PCR on renal tissue for SARS-CoV2 was negative in all 5 biopsies and positive at low level in 6 out of 32 additional kidneys from 16 COVID-19 patients analyzed postmortem in our center (**Supplementary Information 1**), consistently with recent reports suggesting that direct infection of the kidney is possible but other mechanisms of kidney damage might be more relevant in COVID-19 associated AKI.^17–19^ The 5 patients suffered from stage 1 to 3 AKI, as defined by the KDIGO classification in consideration of serum creatinine or urine output, and kidney biopsy was obtained 6 to 26 days after AKI onset. One patient presented clinical and histological evidence for thrombotic microangiopathy (**Extended Data Figure 1**), and all patients displayed tubular alterations at different stages after tubular injury and of variable severity, (**Table 1**, **Figure 1a, b**), determining an ideal model to study the early renal response to AKI in critically ill patients, a clinical context in which kidney biopsies are rarely obtained.

**Table 1.** Clinical characteristics of the patients. AKI: acute kidney injury; HN: hypertensive nephropathy; TMA: thrombotic microangiopathy; VV-ECMO veno-venous extracorporeal membrane oxygenation.

|  | Normal range | #1 | #2 | #3 | #4 | #5 |
| --- | --- | --- | --- | --- | --- | --- |
| **Demographics** |  |  |  |  |  |  |
| Gender |  | female | male | female | male | male |
| Age |  | 79 | 57 | 69 | 76 | 69 |
| BMI (kg/m<sup>2</sup>) |  | 34 | 25 | 20 | 28 | 26 |
| **Comorbidities** |  |  |  |  |  |  |
| Hypertension |  | 1 | 0 | 0 | 1 | 0 |
| Diabetes |  | 1 | 0 | 0 | 0 | 0 |
| Chronic obstructive pulmonary disease |  | 0 | 0 | 0 | 0 | 0 |
| Chronic kidney disease |  | 0 | 0 | 0 | 0 | 0 |
| History of cancer |  | 1 | 1 | 0 | 0 | 0 |
| **Respiratory involvement, complications and severity scores** |  |  |  |  |  |  |
| Mechanical ventilation (days) |  | 10 | 2 | 21 | 17 | 26 |
| VV-ECMO |  | 0 | 0 | 0 | 0 | 1 |
| Septic shock |  | 0 | 1 | 0 | 0 | 1 |
| Deep vein thrombosis |  | 0 | 0 | 0 | 0 | 1 |
| APACHE II score |  | 19 | 33 | 8 | 30 | 15 |
| SAPS II score |  | 82 | 78 | 40 | 70 | 46 |
| SOFA score |  | 7 | 8 | 7 | 3 | 9 |
| **Lab values at the day of kidney biopsy** |  |  |  |  |  |  |
| Hemoglobin (g/L) | 140-180 | 80 | 67 | 74 | 117 | 49 |
| Leukocytes (G/L) | 4-11 | 9.3 | 12 | 10.4 | 8.5 | 24.9 |
| Platelets (G/L) | 150-350 | 326 | 13 | 191 | 262 | 166 |
| Neutrophils (G/L) | 1.5-7.5 | 8.18 (88%) | 10.08 (84%) | 9.05 (87%) | 7.57 (89%) | 22.41 (90%) |
| Lymphocytes (G/L) | 1-4.5 | 0.09 (1%) | 0.36 (3%) | 0.94 (9%) | 0.34 (4%) | 0.87 (3.5%) |
| D-Dimer (ng/mL) | 45-500 | 5416 | 8810 | 795 | 2459 | 9865 |
| LDH (U/L) | 87-210 | 339 | 463 | 360 | 257 | 572 |
| C3 (g/L) | 0.66-1.35 | 1.75 | 1.48 | 1.3 | 1.78 | 0.8 |
| C4 (g/L) | 0.08-0.34 | 0.28 | 0.26 | 0.19 | 0.27 | 0.2 |
| TNFα (pg/mL) | <4 | 17.96 | 5.6 | 6.16 | 1.42 | 3.2 |
| IL6 (pg/mL) | <1.5 | 73.9 | 1190.5 | 474.1 | 3.76 | 6.8 |
| IL8 (pg/mL) | <90 | 122.5 | 84.2 | 290 | 45.91 | 32.5 |
| IL10 (pg/mL) | <1 | 2.5 | 5.9 | 3.33 | <1 | 3.8 |
| MCP1 (pg/mL) | 50-260 | 1539 | 1243.2 | 1715 | 302 | 115.2 |
| **Kidney function** |  |  |  |  |  |  |
| Baseline serum creatinine (μmol/l) |  | 150 | 95 | 66 | 76 | 80 |
| Peak serum creatinine (μmol/l) |  | 351 | 586 | 80 | 96 | 109 |
| Serum creatinine at sampling (μmol/l) |  | 175 | 219 | 61 | 62 | 39 |
| AKI stage (KDIGO) |  | 2 | 3 | 2 | 2 | 1 |
| AKI criterium |  | creatinine | creatinine | oliguria | oliguria | oliguria |
| Days between AKI onset and biopsy |  | 6 | 9 | 18 | 15 | 25 |
| Renal replacement therapy |  | 0 | 1 | 0 | 0 | 0 |
| **Urinalysis at the day of biopsy** |  |  |  |  |  |  |
| Protein (g/L) |  | 0.56 | 1.21 | 1.26 | 0.44 | 0.13 |
| Albumin (mg/L) (<10) |  | 65 | 82 | 42 | 12 | 11 |
| Creatinine (mmol/l) |  | 1.9 | 2.3 | 3.9 | 5.3 | 3.3 |
| Phopshate (mmol/l) |  | 10.5 | 18.4 | 23 | 50 | 18.2 |
| Magnesium (mmol/l) |  | 2.33 | 2.41 | 5.18 | 6.37 | 2.22 |
| **Histological findings** |  |  |  |  |  |  |
| Tubule |  | Severe tubular atrophy | Moderate acute tubular lesion and atrophy | Moderate acute tubular lesion | Slight tubular atrophy | Moderate tubular atrophy |
| Interstitial fibrosis |  | Slight | Slight | Slight | Moderate | Slight |
| Interstitial inflammation |  | Focal | Absent | Focal | Absent | Absent |
| Pathology others |  |  | TMA |  | HN |  |

**Figure 1.**
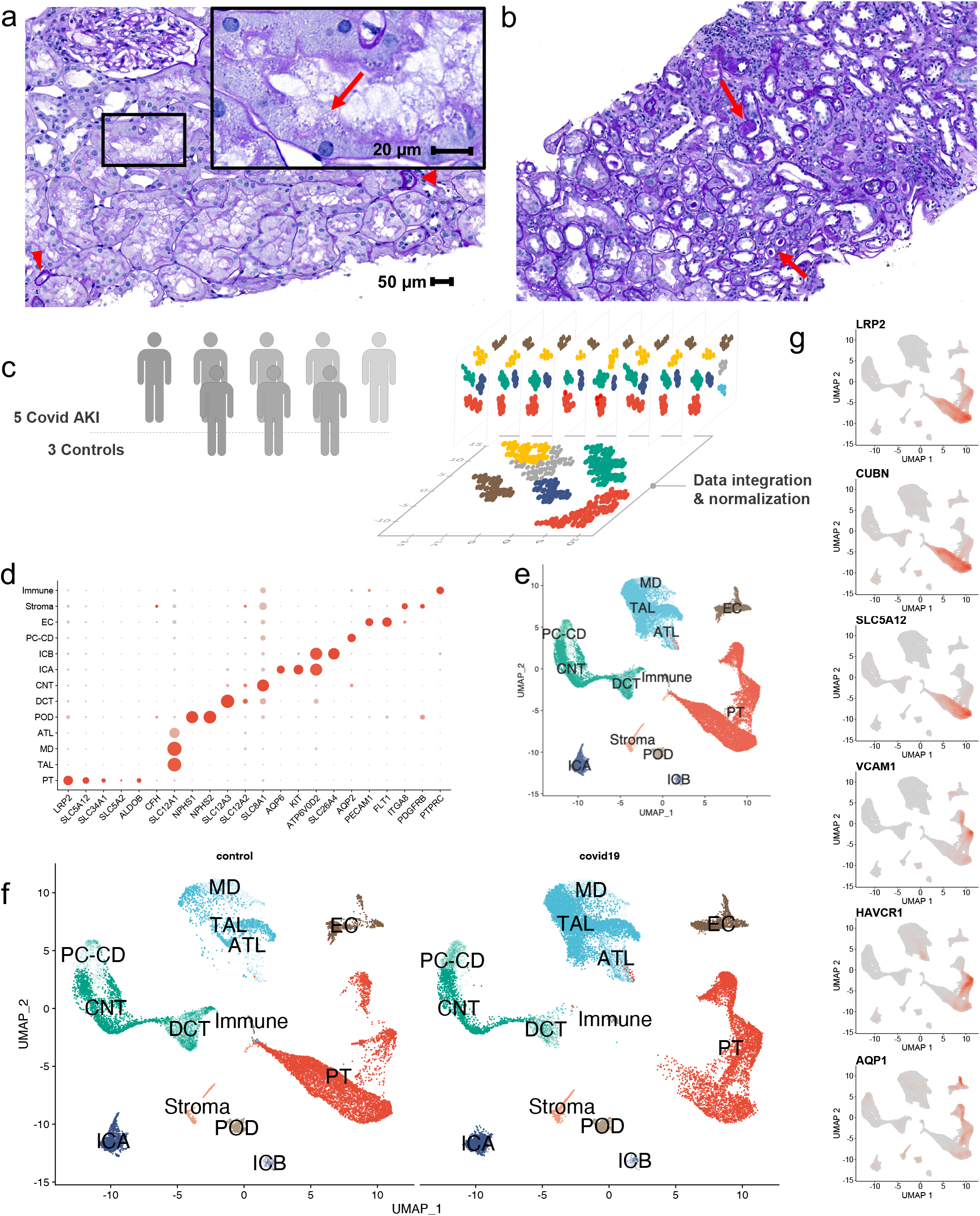
a) Histopathology of a representative renal cortical area showing diffuse acute tubular lesions characterized by non-isometric sloughing of tubular epithelial cell and brush border loss (boxed area). Sparse foci of atrophic tubules are noted (red arrows heads). Red arrow indicates complete loss of the brush border. Periodic Acid-Schiff stain, b) Representative deeper cortical section showing advanced chronic lesion with moderate inflammation and interstitial fibrosis with atrophic tubules and tubular cast formation (red arrows). Rare foci of acute tubular lesion are noted. Periodic Acid-Schiff stain, c) Schematic of the data integration strategy, d) Dotplot of cell cluster marker genes identified in Covid-19 AKI and control samples (dot size indicates the percentage of positive cells and color indicated relative expression), e-f) UMAP representation of the 13 cell types identified by unsupervised clustering of the whole dataset (e) and split in the control and Covid-19 groups (f), g) scatterplot from the UMAP projection showing the gene-weighted density of genes of interest. PT, proximal tubule; ATL, thin ascending limb of the loop of Henle; TAL, thick ascending limb of the loop of Henle; POD, podocytes; MD, macula densa; DCT, distal convoluted tubule; CNT, connecting tubule; ICA, type A intercalated cells of the collecting duct; ICB, type B intercalated cells of the collecting duct; PC, principal cells; CD, collecting Duct; EC, endothelial cells.

The snRNAseq data were first integrated with 3 controls from available datasets generated with similar tissue processing and single cell technology for data control and validation (**Figure 1c**).^5^ After data processing and quality control, we obtained 35,299 single cell transcriptomes (20,165 from COVID19 patients and 15,134 from controls (**Extented Data Figure 2**). Based on established cell type markers we identified all expected kidney cell populations (**Figure 1d, e and Extended Data Figure 3, Supplementary information 2**). The number of immune cells detected by snRNAseq was very low, reflecting the absence of prominent inflammatory infiltrates observed by conventional histology. The most prominent finding in the snRNAseq analyses was in the proximal tubule (PT) compartment (marked by LRP2 and CUBN) (**Figure 1 f, g**), consistently with the pathological findings observed by conventional histology (**Figure 1a, b**). We found abnormal PT cells in all samples obtained from COVID-19 patients, including patients with minimal histological alterations.

In the focused analysis on the PT, we found a subset of cells displaying the classical transcriptional profile of mature PT cells and a large heterogenous cluster of abnormal cells (**Figure 2a, b**). The general comparison between mature and abnormal PT cells highlighted the reduction of genes associated with renal tubule functions, such as electrolyte transport and fatty acid metabolism, and activation of NOTCH signaling (**Figure 2c**). A subset of those undifferentiated PT cells displayed classical markers of tubular injury, such as HAVCR1 and VCAM1 (**Figure 2d**). Secondary validation by immunohistochemistry indicated that cells marked by HAVCR1 and VCAM1 corresponded to the undifferentiated, flattened tubular cells, typically observed after tubular injury (**Figure 2e, f and Extended Data Figure 4**).^20^ The general characterization of the undifferentiated PT cells highlighted an altered cell metabolism, with a switch from fatty acid oxidation and gluconeogenesis to glycolysis, a shift from de novo to salvage pathway in NAD biosynthesis and evidence for altered mitochondrial biogenesis (**Figure 2g**).^21–23^ Moreover, undifferentiated PT cells displayed higher extra-cellular matrix scores, consistent with their potential role in the mechanisms determining the transition to chronic kidney disease (**Figure 2g**), as previously postulated^2,6^ and in line with the observation of fibrotic tissue surrounding HAVCR1 marked cells in the patients with a prolonged delay between AKI onset and kidney biopsy (**Figure 2h-i)**.

**Figure 2.**
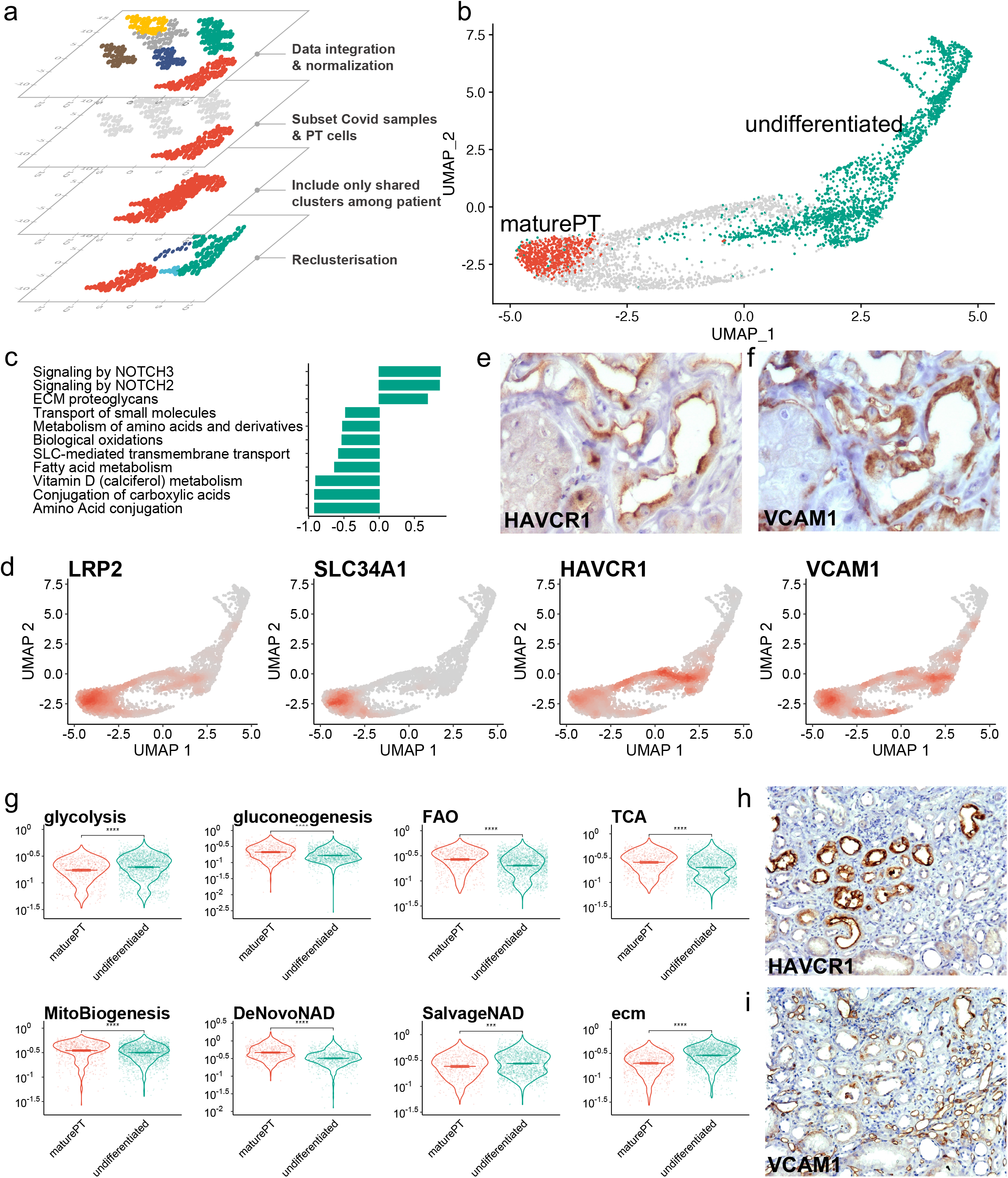
a) Representative schematic of the clustering strategy to focus analyses on PT cells from Covid-19-AKI samples, b) Scatterplot showing the projection of two cell types identified by supervised clustering of the PT dataset projected on the UMAP, c) Barplots showing the enrichment score of PT cells compares to undifferentiated cells, calculated by Gene Set Enrichment Analysis using Reactome pathways database (positive enrichment means an enrichment in undifferentiated cluster), d) Scatterplot from the UMAP projection showing the gene-weighted density of genes of LRP2, SLC34A1, HAVCR1 and VCAM1, e,f) Representative immunostainings of HAVCR1 (e) and VCAM-1 (f) proteins in atrophic tubules, g) Violin plots showing the activity of the the following pathways: glycolysis, gluconeogenesis, Fatty Acid Oxidation (FAO), Tricarboxylic Acid Cycle (TCA), Mitochondrial biogenesis (MitoBiogenesis), NAD synthesis from de novo pathway (DeNovoNAD), NAD synthesis from salvage pathway (SalvageNAD) and ExtraCellularMatrix (ecm), in mature PT and undifferentiated cells, h,i) Representative immunostainings of HAVCR1 (h) and VCAM-1 (i) proteins in fibrotic area, *** p < 0.001 **** p < 0.0001.

Normal and undifferentiated PT cell clusters were connected by two cell state transitions (trajectory #1 and #2 in **Figure 3a, Supplementary information 3**). Trajectory #1 was more prominent in patient with more severe AKI (i.e. in patients with relevant increase in serum creatinine) and higher inflammatory cytokine levels (**Extended Data Figure 5**). The gene expression profile characterizing cells in trajectory #1 was reminiscent of the very early response to AKI previously characterized in kidney biopsies performed after organ reperfusion in the setting of transplantation^13^ (e.g. increase in genes associated with EGFR, VEGF), but this was not the case for cells in trajectory #2 (**Extended Data Figure 6**). Therefore, we hypothesized that trajectory #1 would correspond to the early tubular response to injury, leading to cell de-differentiation (and likely cell death in a subset of cells), whereas trajectory #2 would represent the processes of cell de- and re-differentiation along the mechanisms of cell plasticity in tubule repair previously characterized in mice (**Figure 3b**).^7,10^ We used a mouse model of ischemia-reperfusion injury (IRI) to verify the hypothesis and compare this processes across species. We extracted from snRNAseq data the transcriptional profiles of PT cells in the first hours/days after injury to study the early response to injury (**Figure 3c, Extended Data Figure 7**). Moreover, to track the PT re-differentiation process, we injected 5-ethyny-2’-deoxyuridine (EDU) 48h after moderate ischemia-reperfusion injury and we sorted EDU+ nuclei 48h and 26 days later (**Figure 3d, Extended Data Figure 8, Supplementary information 4**). Reparative cells proliferate and integrate EDU in the early phase after injury and maintain this label in the following days.^24^ EDU+ cells were in a dedifferentiated cell state 96h but recovered to a fully differentiated phenotype at day 28 after injury.^21^ The comparison between this controlled injury/repair process and the human dataset highlighted similar gene expression transitions in the early and in the reparative phase across species (**Figures 3e-g**). For a better characterization the underlying biological processes, we identified genes following similar gene change patterns along the pseudotime and we performed gene enrichment analysis in the corresponding gene sets. The early injury trajectory was characterized by the reduction first of genes related to cell metabolism and then to ion transport, in parallel we observed an increase in genes associated with cell death, toll-like receptor signaling and cell adhesion molecules (**Extended Data Figure 9**). Along the re-differentiation transition PT cells re-acquired their classical metabolic and transporter functions and progressively lost genes involved in cell mitosis, adhesion and motility, and genes associated with EGF, PDGF and AGR signaling; we also found a set of genes associated with epithelial morphogenesis showing a transient increase along the trajectory (**Extended Data Figure 10**). Thus, the early response to injury and the process of tubule repair by dedifferentiation and redifferentiation of mature PT cells similarly occurs in mouse and human.

**Figure 3.**
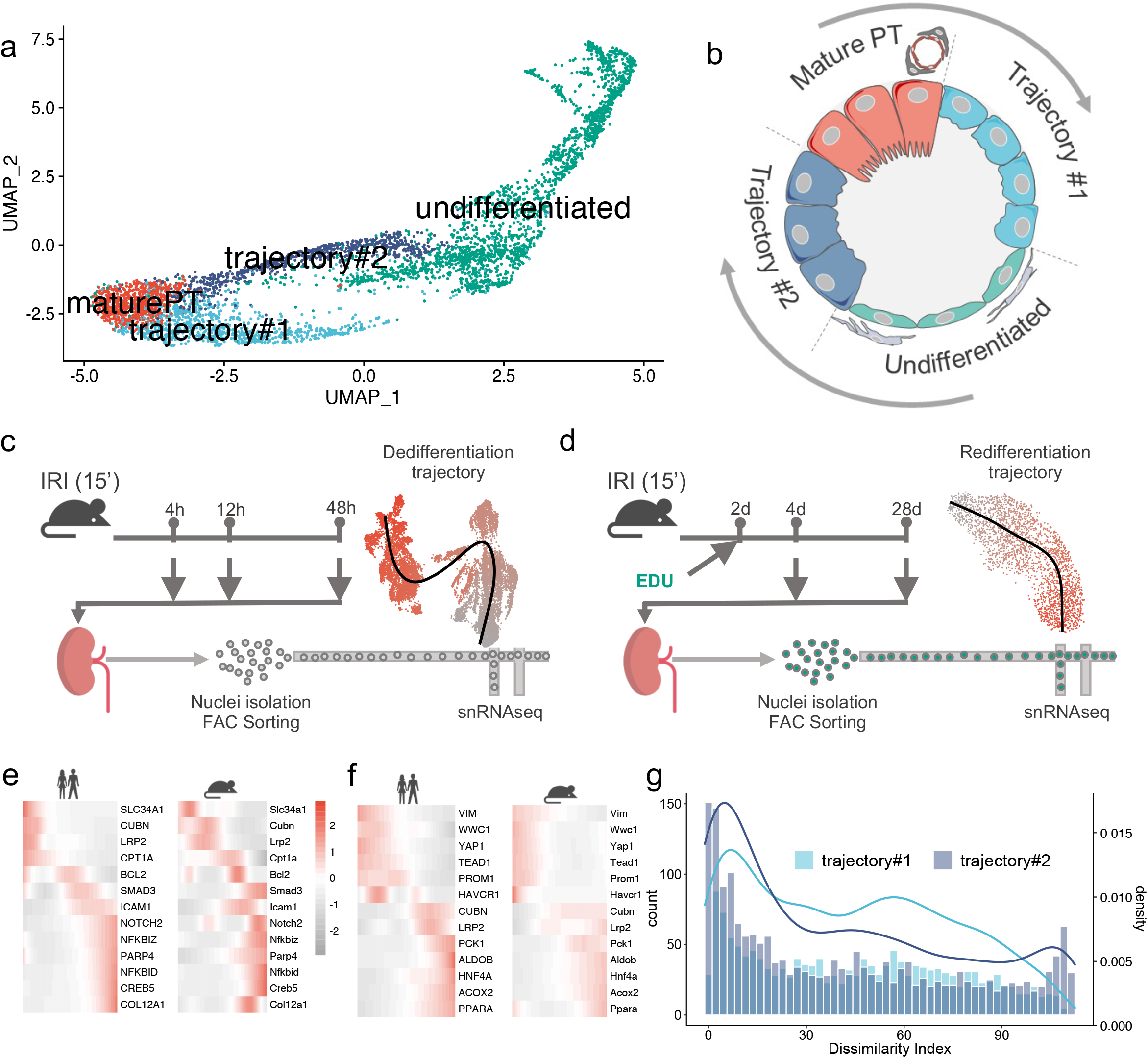
a) Scatterplot showing the projection of the four cell types identified by supervised clustering of the PT dataset projected on the UMAP, b) Schematic model of the injury and repair processes undergoing in PT cells, c,d) schematic diagram of the experimental procedures to study the dedifferentiation (c) and redifferentiation (d) processes: mice underwent ischemia reperfusion induced acute kidney injury (with an EDU injection at day2 for the redifferentiation process) and were sacrificed at different timepoints (4h, 12h, 48h, 4 days and 28 days). The kidneys were harvested, the nuclei isolated by FAC sorting via DAPI staining and sequenced at single cell resolution. After data integration and clusterisation, the UMAP reductions were used as an input for pseudotime inference by slingshot (black line) with color indicating the timeline of the pseudotime, grey beeing pseudotime 0 and dark red the latest pseudotime, e,f) Heatmap showing the expression pattern over pseudotime of selected AKI related genes in man (left) and mouse (right), for the cell differentiation (e) and redifferentiation (f) processes, g) histogram and density curves showing the distribution of the dissimilarity index, calculated for each couple of genes, shared by the Covid19 and the mice model and for each trajectory#1 and trajectory#2.

Undifferentiated PT cells displayed a higher level of cell heterogeneity in man than in mouse, with the presence of additional cell clusters marked by PROM1 and CD24 in the human dataset, reminiscent of multipotent progenitor cells previously described in adult human kidney (**Figure 4a**).^8–10^ PROM1 expression was mutually exclusive with markers of tubular injury, as assessed by snRNAseq and validated by immunohistochemistry, whereas we observed a partial overlap between PROM1 and PAX8 (**Figure 4b, Extended Data Figure 11**). In fact, pseudotime analysis revealed a sequential expression of genes involved in stem and progenitor cell biology, starting from a population enriched for LGR5, KRT7, CD24 and other progenitor cell markers (**Figure 4c**).^8,25^ However, PROM1 positive cells identified by immunohistochemistry did not form defined clusters but were broadly distributed in tubules undergoing repair (**Figure 4b**). This finding was consistent with the activation of a reparative response determining the transition to a progenitor-like transcriptional profile and not with the expansion of predefined progenitor cells in the adult human kidney. To comprehensively analyze the cell state transition from this progenitor-like cell state to mature PT cells we identified genes following similar gene change patterns along the pseudotime and we performed gene enrichment analysis in the corresponding gene sets. The cells progressively lost the expression of genes associated with kidney development (e.g. EYA1, FGF1, **Figure 4e, Supplementary Information 5**). We found an intermediate state related to cell proliferation and change in cell structure and organelle organization (**Figure 4d,f, Supplementary Information 5**), followed by the expression of genes related to carbohydrate, amino acid and lipid metabolism and transmembrane transport, determining the differentiation into mature PT cells (**Figure 4g, Supplementary Information 5**). A specific analysis on the main pathways involved in this process revealed the early activation of WNT, followed by NOTCH, HIPPO and calcium signaling, with an increase in HNF4 activity in the last cell maturation steps (**Figure 4h**).

**Figure 4.**
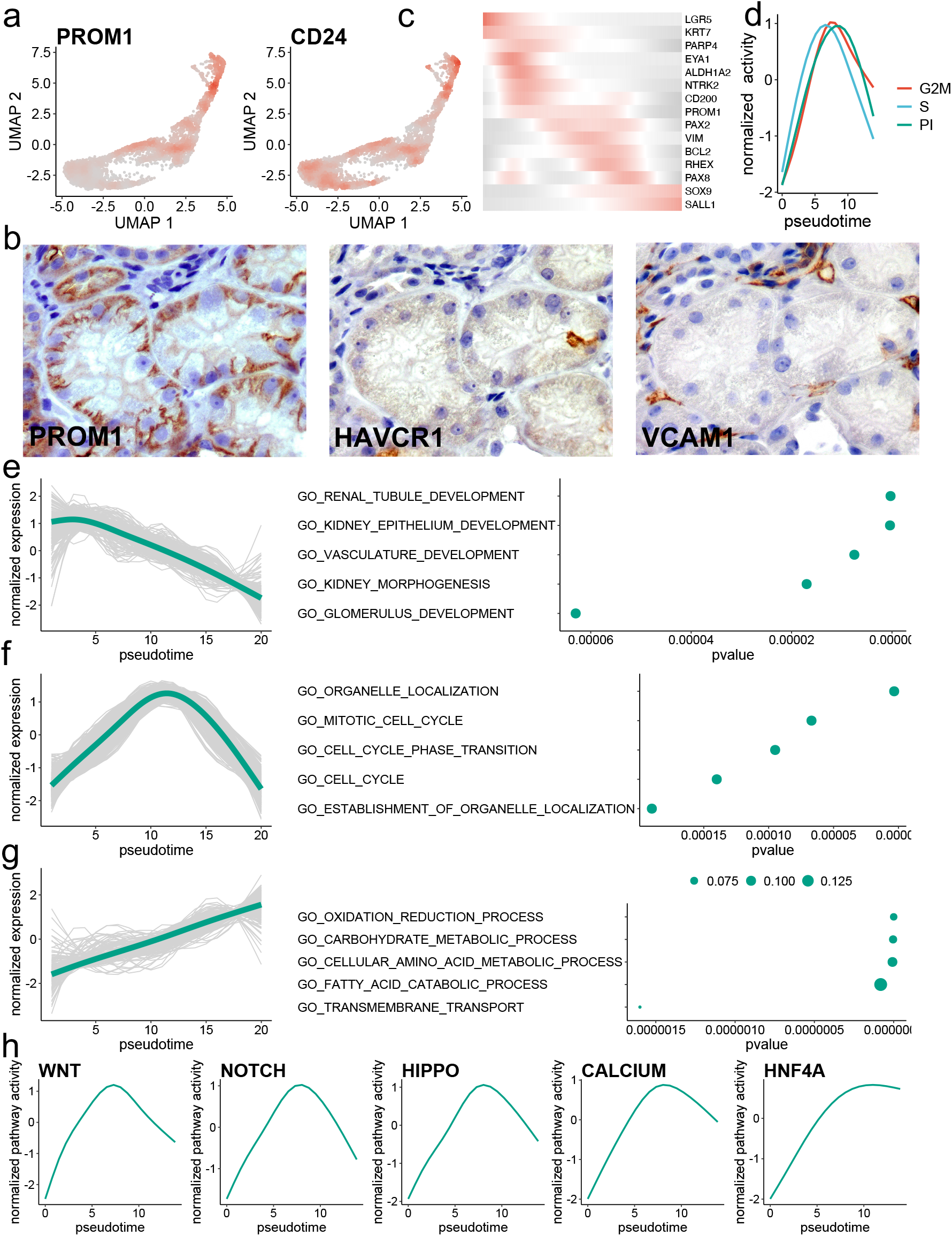
a) Scatterplot from the UMAP projection showing the gene-weighted density of genes of PROM1 and CD24, b) Representative immunostainings for PROM1 (left panel), HAVCR1 (middle panel) and VCAM1 (right panel) proteins in tubules (controls are shown in **Extended Data Figure 4**), c) Heatmap showing the expression pattern over pseudotime of kidney progenitor genes, from undifferentiated cluster to mature PT, d) score activity along pseudotime for G2M-phase (red), S-phase (green) and Proliferation Index (PI, blue), e-g) pathway enrichment using genes with a similar expression pattern along time as an input, left panels showing the individual gene expression along time in grey and the generalized additive smooth model in green, middle panels showed a selection of 5 or the most associated pathways and right panels the related P-values and h) normalized pathway activity along pseudotime.

Thus, by integrating snRNAseq data from several kidney biopsies in the first weeks after AKI in critically ill patients with COVID19 we generated the first map of PT injury and repair in human. Early injury response and the fundamental processes of tubule cell plasticity were similar in mouse and man. However, the activation of a reparative response starting from a progenitor-like cell state and partially recapitulating nephrogenesis has not been observed in the mouse but is likely to involve previously identified cell types.^9,25^ An intermediate cell state characterized by loss of PT functions and altered metabolic properties emerged as a common step in the process of tubule injury and along the tubule repair process with indirect evidence suggesting a potential role of those cells in promoting fibrosis. The COVID19 pandemics determined a unique clinical setting to study kidney repair in severely ill patients and to characterize for the first time the fundamental cellular mechanisms of tubule repair in the early phase after AKI at single cell resolution.

## Acknowledgments

We thank Andrea Alimonti for supporting the project and Lorenzo Ruinelli for data management support.

---

**Extended Data Figure 1.**
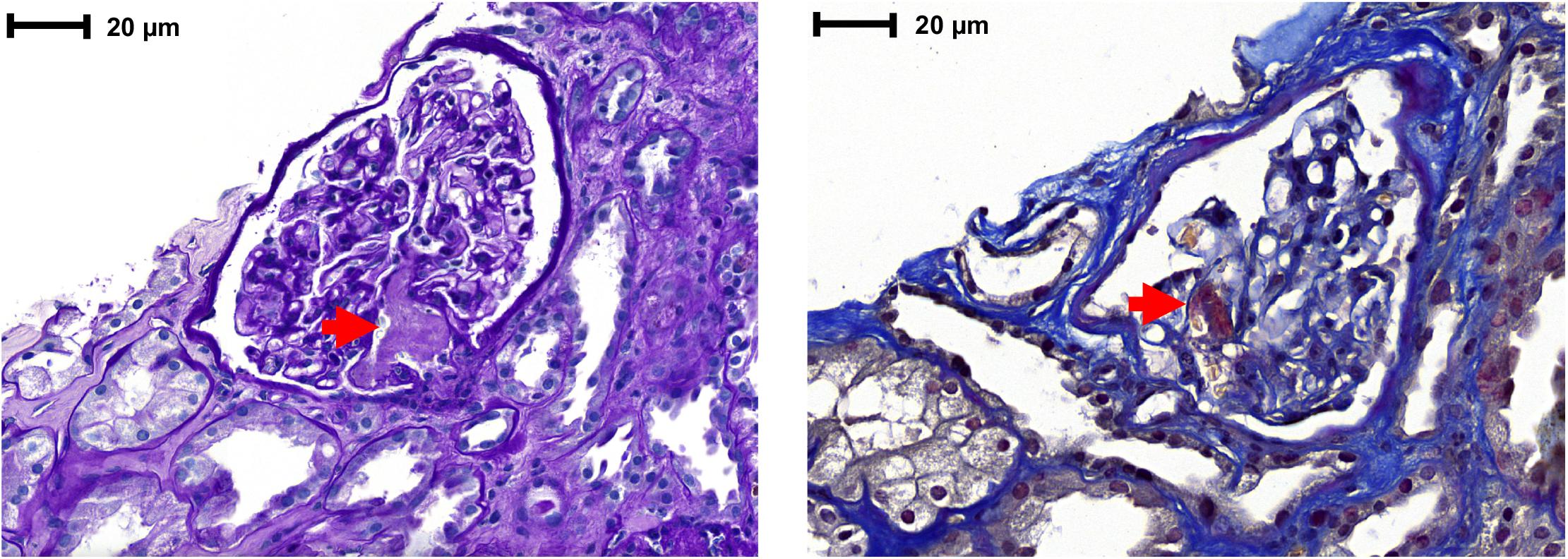
Representative illustration of a renal thrombotic microangiopathy observed in patient #2, characterized by intracapillary fibrinous thrombi and subsequent ischemic alteration with glomerular retraction and non-isometric sloughing of tubular epithelial cell. Red arrows indicated fibrinous thrombus. Periodic Acid-Schiff (left panel) and Masson’s trichrome (right panel) stains.

**Extended Data Figure 2.**
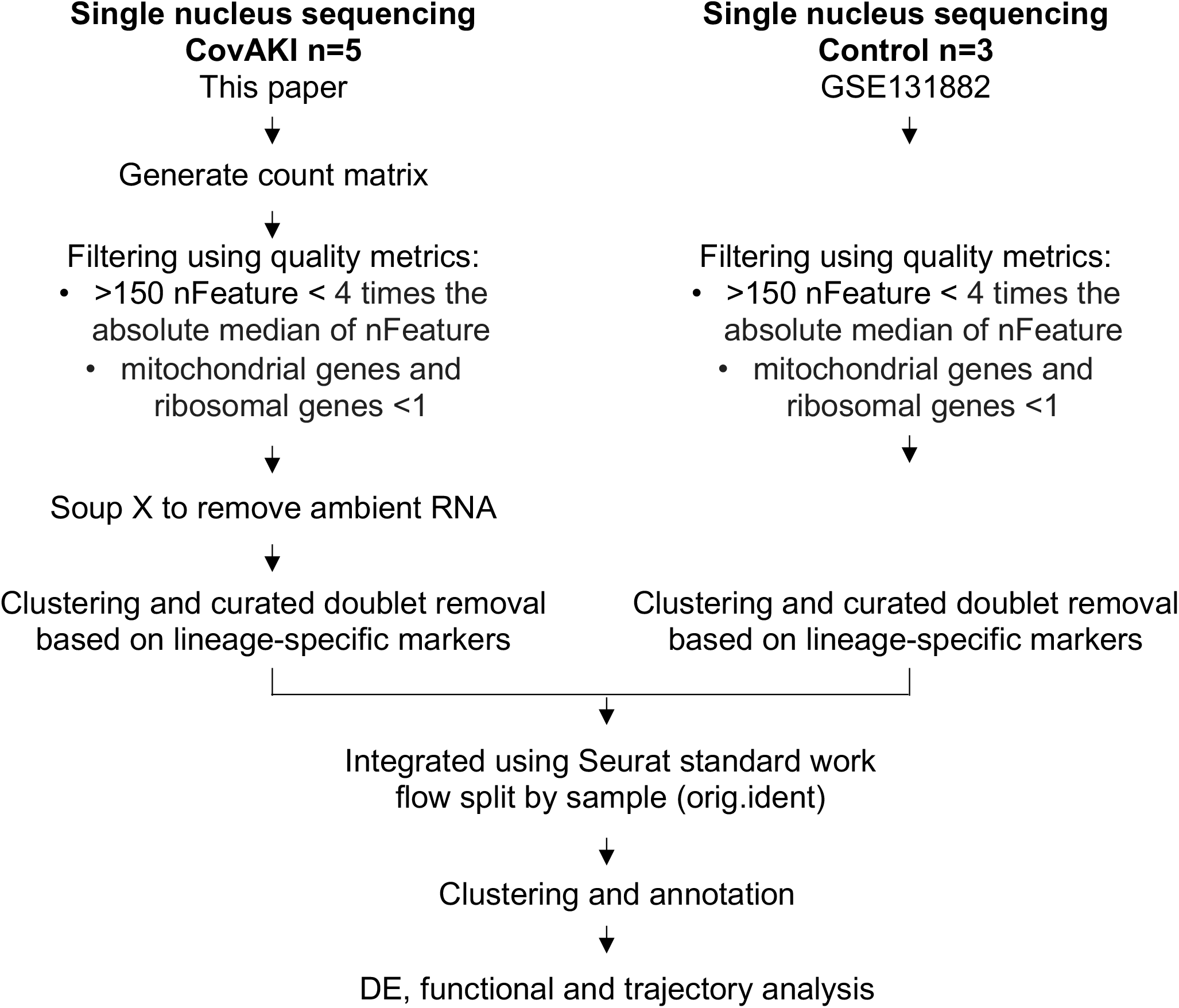
Summary of the data integration strategy

**Extended Data Figure 3.**
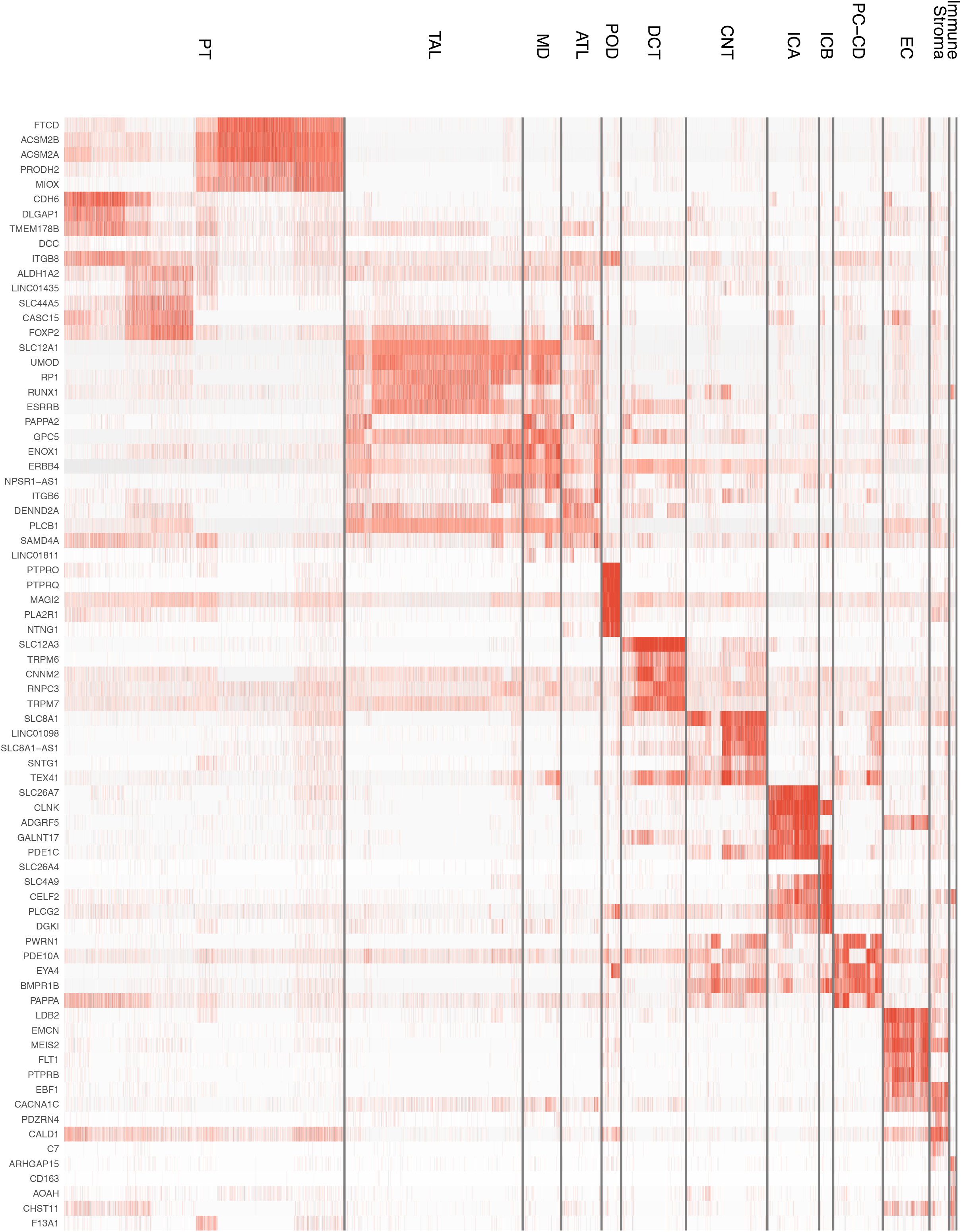
Heatmap showing the expression of the top 5 genes for each cell cluster

**Extended Data Figure 4.**
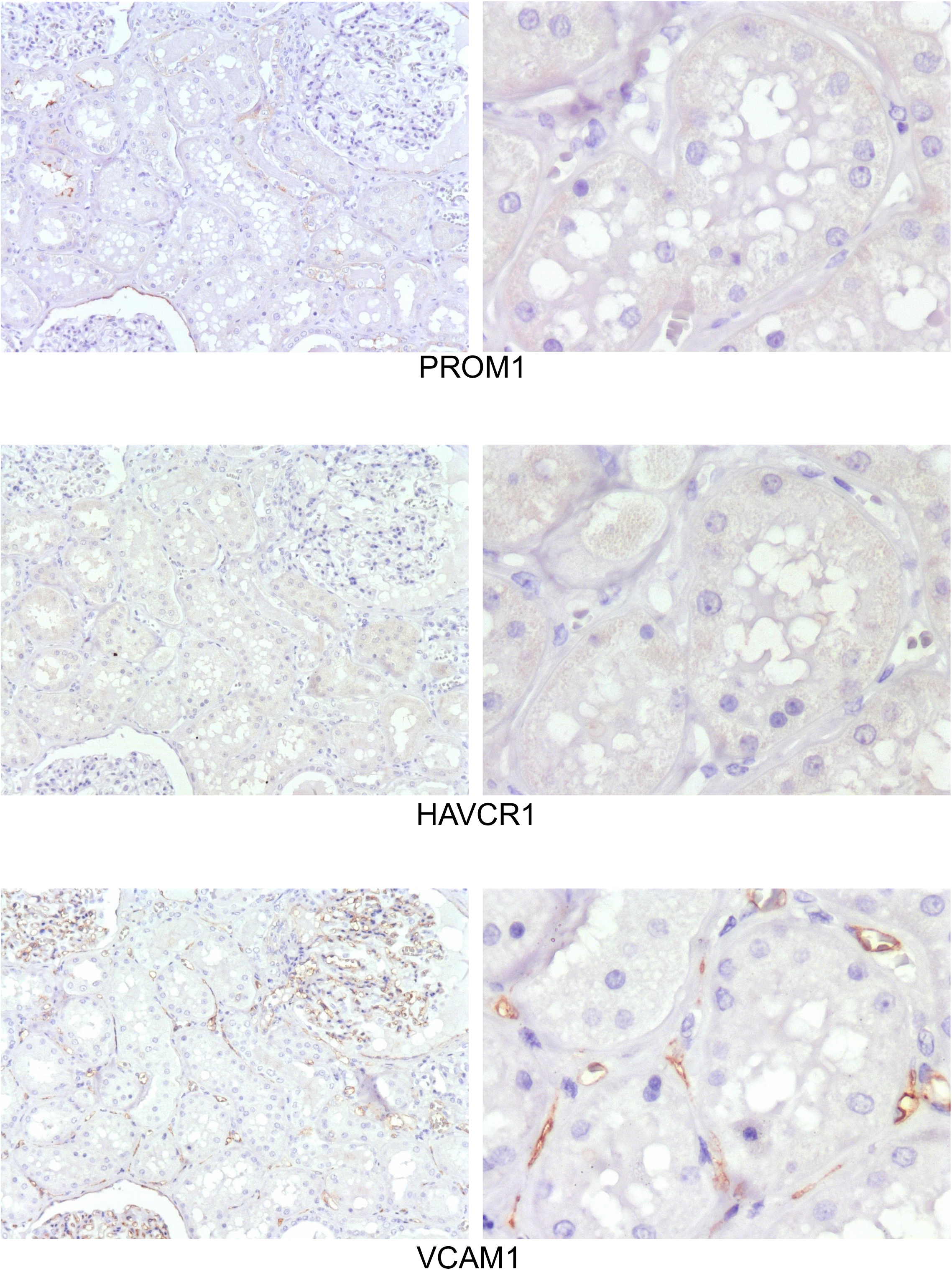
Representative immunostaining of PROM1, HAVCR1 and VCAM1 proteins in normal kidney.

**Extended Data Figure 5.**
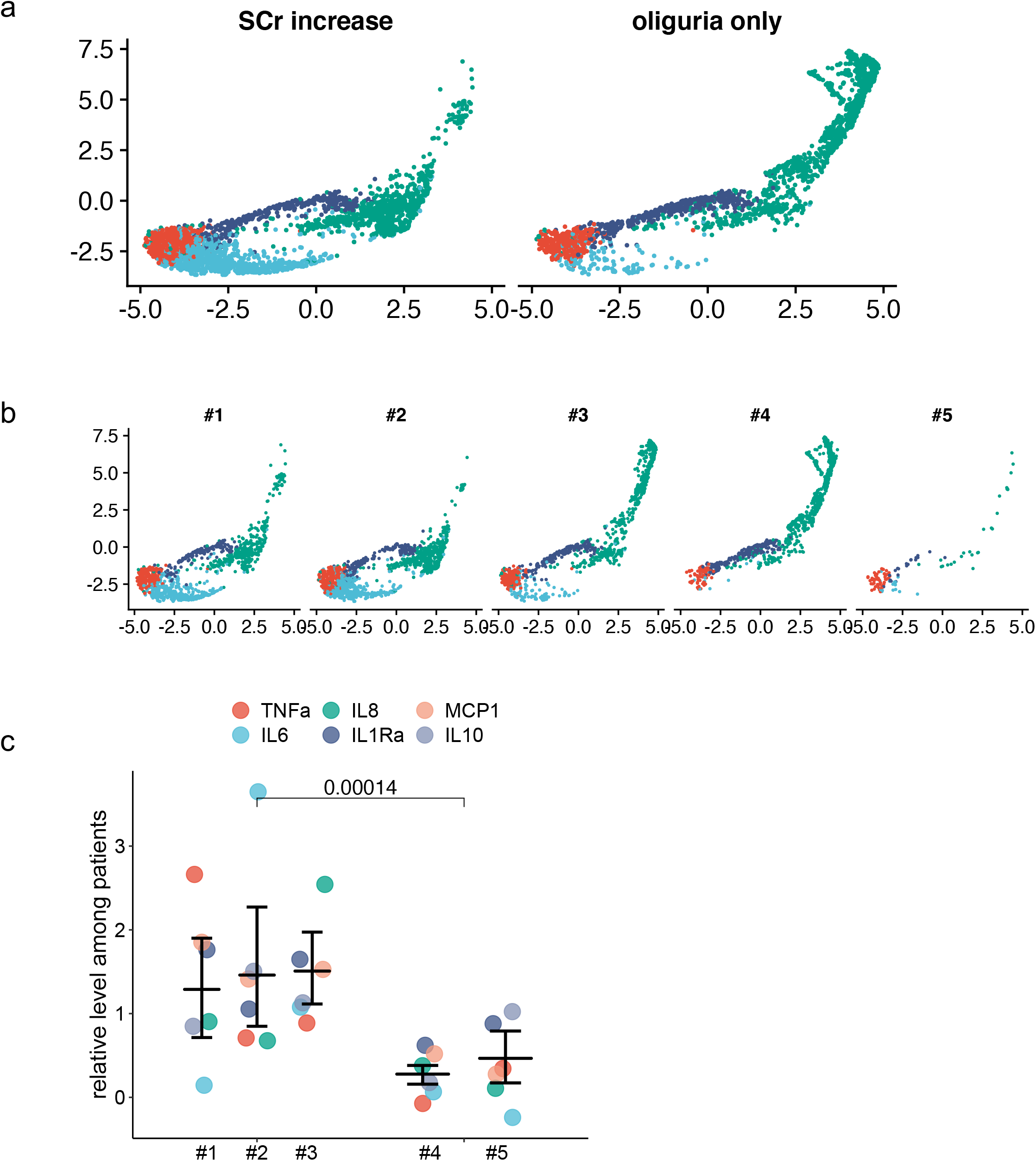
a,b) Scatterplot showing the projection 4 cell states shown in figure 3a identified by unsupervised clustering of the PT dataset projected on the UMAP and split by AKI definition (a) or patient (b) and c) serum level of interleukins in the five included patients. The interleukins level was normalized among patient for each interleukine.

**Extended Data Figure 6.**
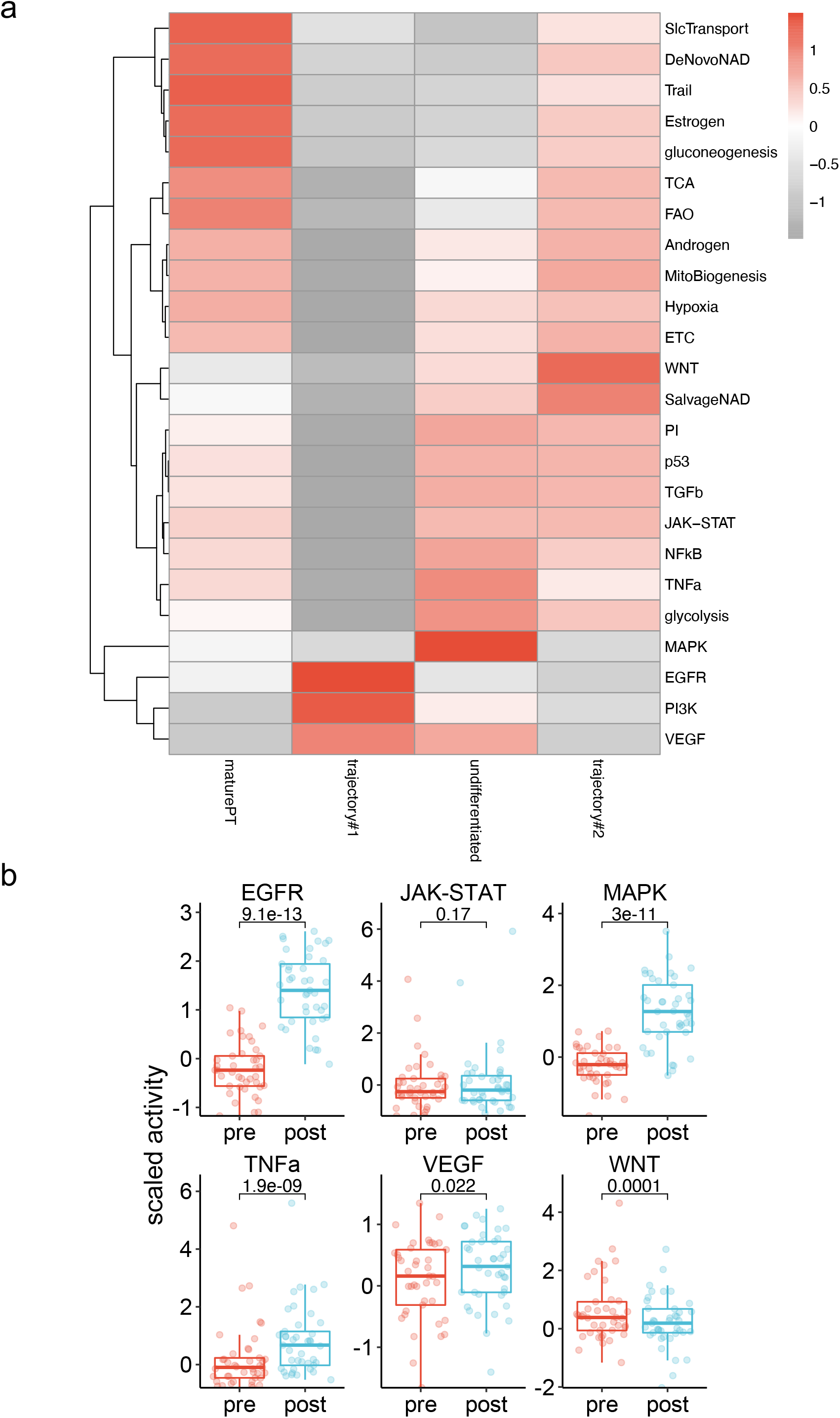
Comparisons of pathways activity in two human datasets, a) Heatmap showing pathways activity in every cell population defined in the PT cells from the Covid19 patients, b) boxplots displaying the pathways activity calculated by Progeny in allograft kidney recipients’ biopsies collected before transplantation or after reperfusion.

**Extended Data Figure 7.**
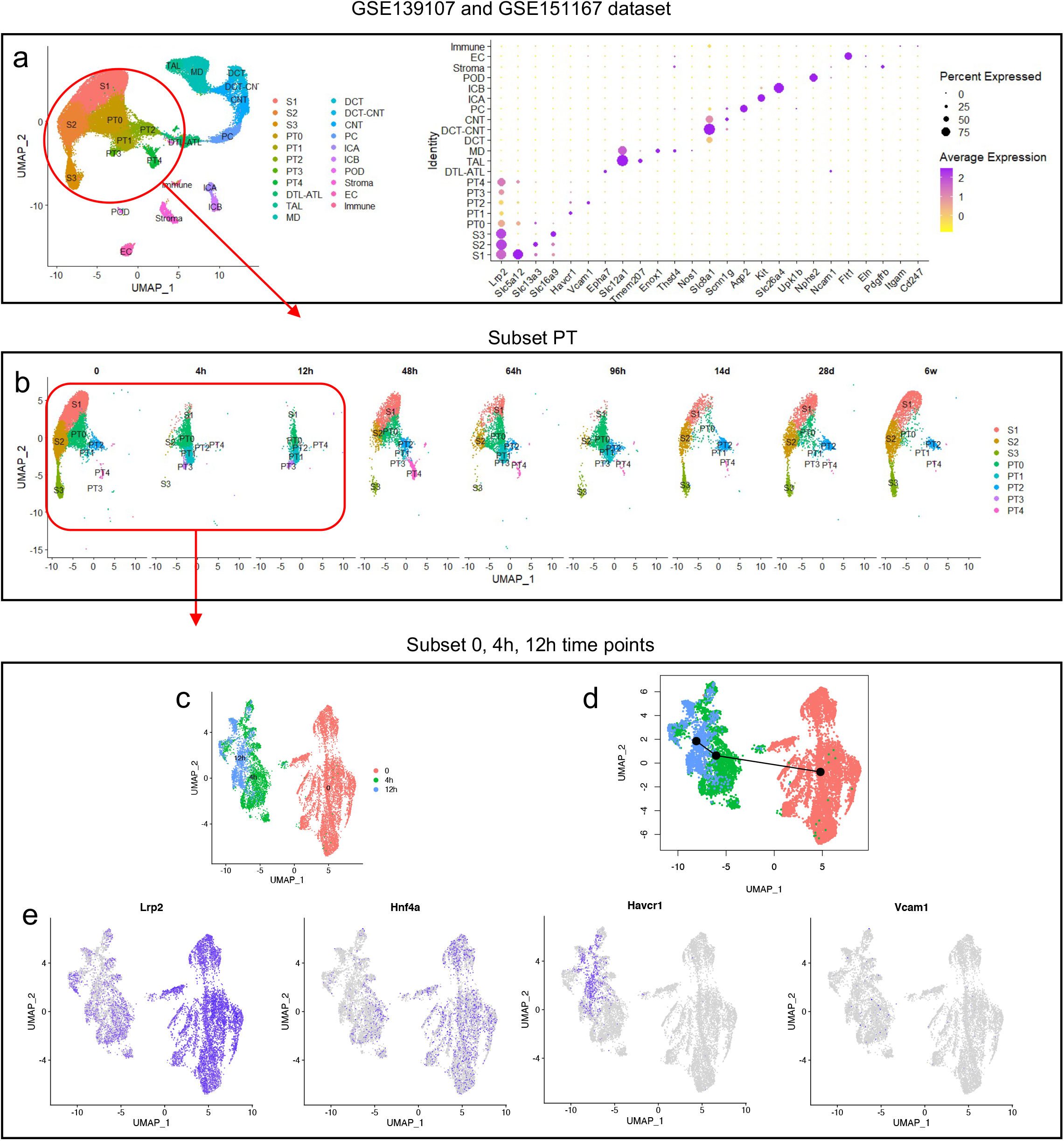
Data analysis pipeline of single nucleus RNA seq data of mouse IRI kidney, a) UMAP plot and dot plot of all integrated datasets (GSE139107, GSE151167) show the identification of the different cellular component of the kidney according to standard markers of renal cell types. PT cells have been selected. PT, proximal tubule cells; DTL, descending limb of loop of Henle; ATL, thin ascending limb of loop of Henle; TAL, thick ascending limb of loop of Henle; POD, podocytes; MD, macula densa; DCT, distal convoluted tubule; CNT, connecting tubule; ICA, type A intercalated cells of collecting duct; ICB, type B intercalated cells of collecting duct; PC, principle cells; EC, endothelial cells, b) UMAP plot shows the time points included in the datasets and in the red box the time points selected to analyse dedifferentiation after injury (32452 genes x 4822 nuclei), c) UMAP of injured PT, d) Trajectory of dedifferentiation using Slingshot, e) Featureplot of markers of differentiation and injury.

**Extended Data Figure 8.**
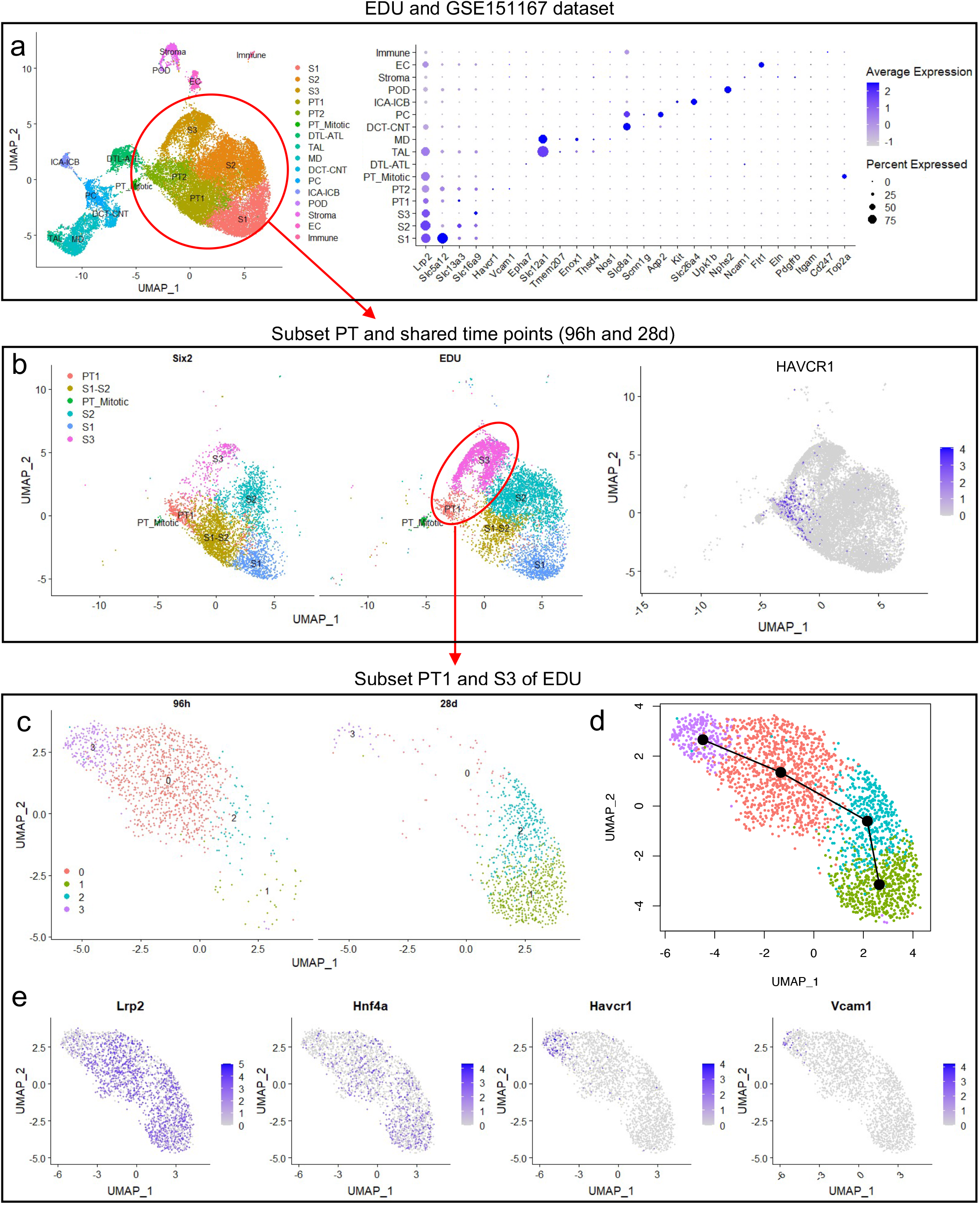
Data analysis pipeline of single nucleus RNAseq data on EDU+ sorted cells obtained and controls (Six2) in the ischemia reperfusion injury (IRI) mouse model, a) EDU and SIX2 dataset integration: EDU+ cells obtained 96 h (n=2 replicates) and 28 days (n=2 replicates) after IRI were merged with sham controls (n=3 replicates) (total 32’452 genes x 29’492 nuclei) and annotated based on established markers, b) PT subset. PT cells and shared time points between datasets (96h and 28d) were selected (32452 genes x 11506 nuclei), c) Subset S3 and PT1 (injured cells) from EDU dataset. Further analyses were focused on those cells because tubular injury was mainly restricted to this part of the nephron in this model (32’452 genes x 2’263 nuclei), d) Trajectory of differentiation using Slingshot. e) Feature plot of markers of differentiation and injury.

**Extended Data Figure 9.**
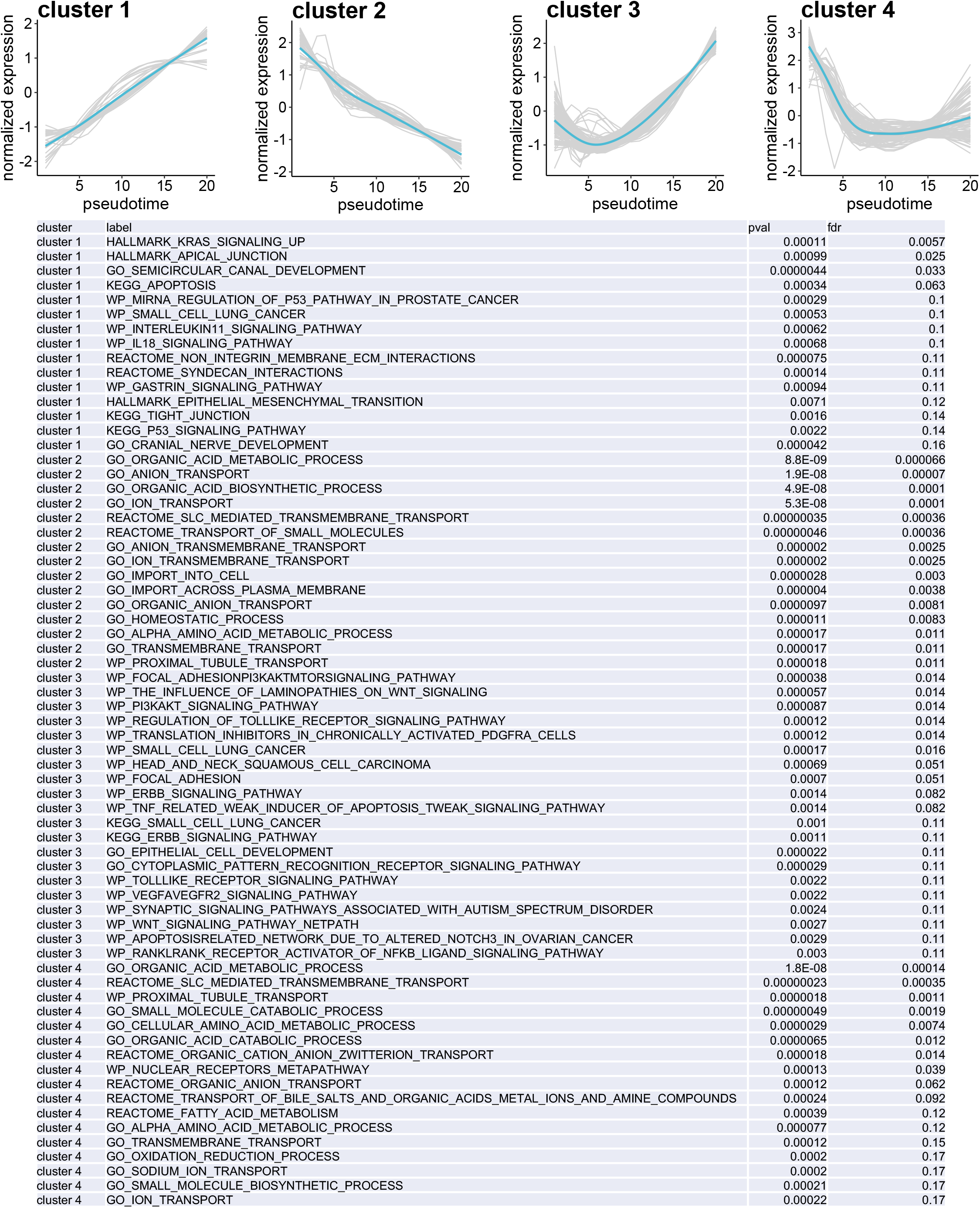
Upper panel shows the individual (in grey) and smooth (in lightblue) gene expression along dedifferentiation trajectory, grouped by clusters of similar pattern and lower table indicates for each cluster of expression pattern, the top15 associated pathways and their relative pvalues and false discovery rates.

**Extended Data Figure 10.**
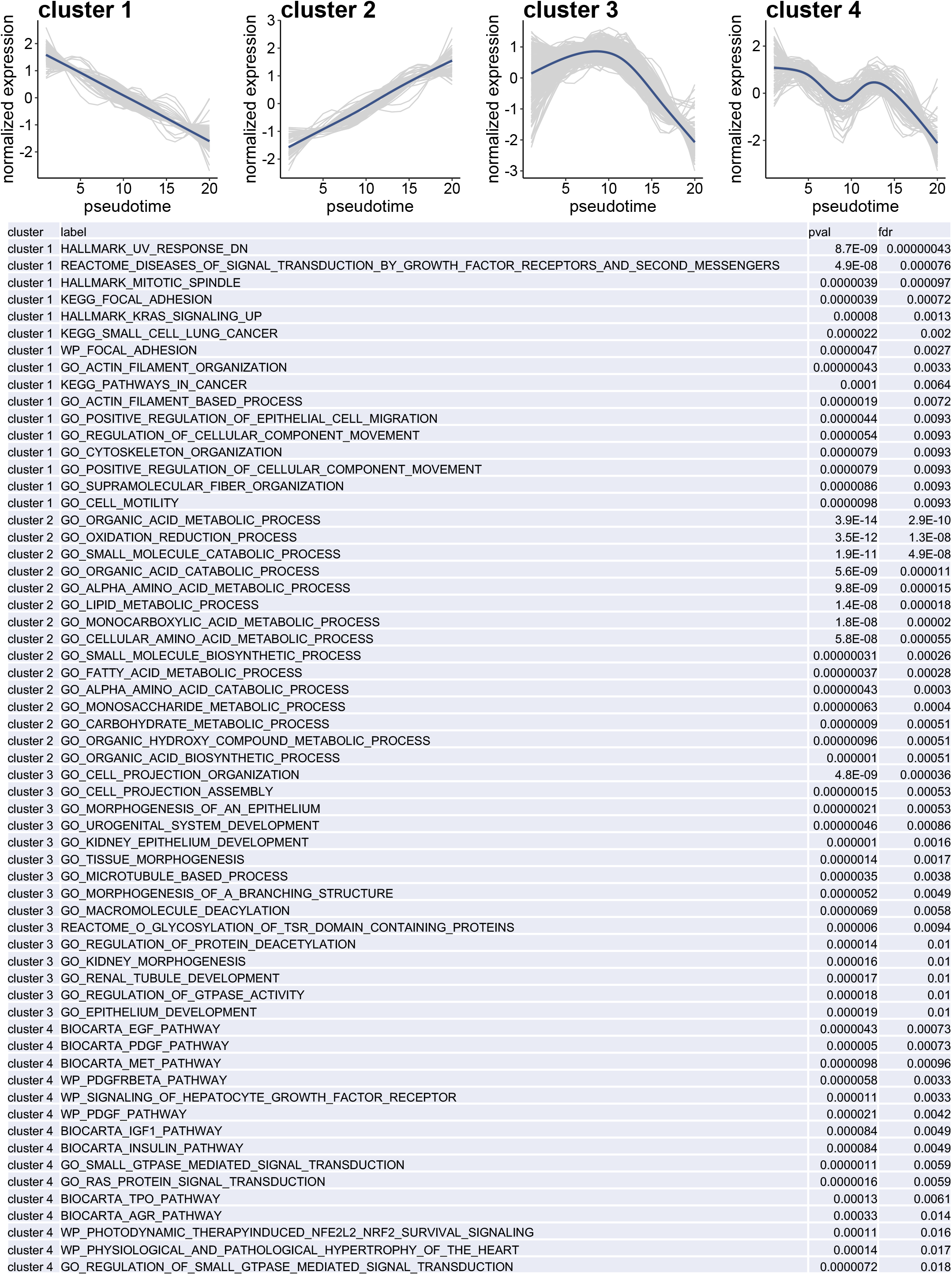
Upper panel shows the individual (in grey) and smooth (in darkblue) gene expression along redifferentiation trajectory, grouped by clusters of similar pattern and lower table indicates for each cluster of expression pattern, the top15 associated pathways and their relative pvalues and false discovery rates.

**Extended Data Figure 11.**
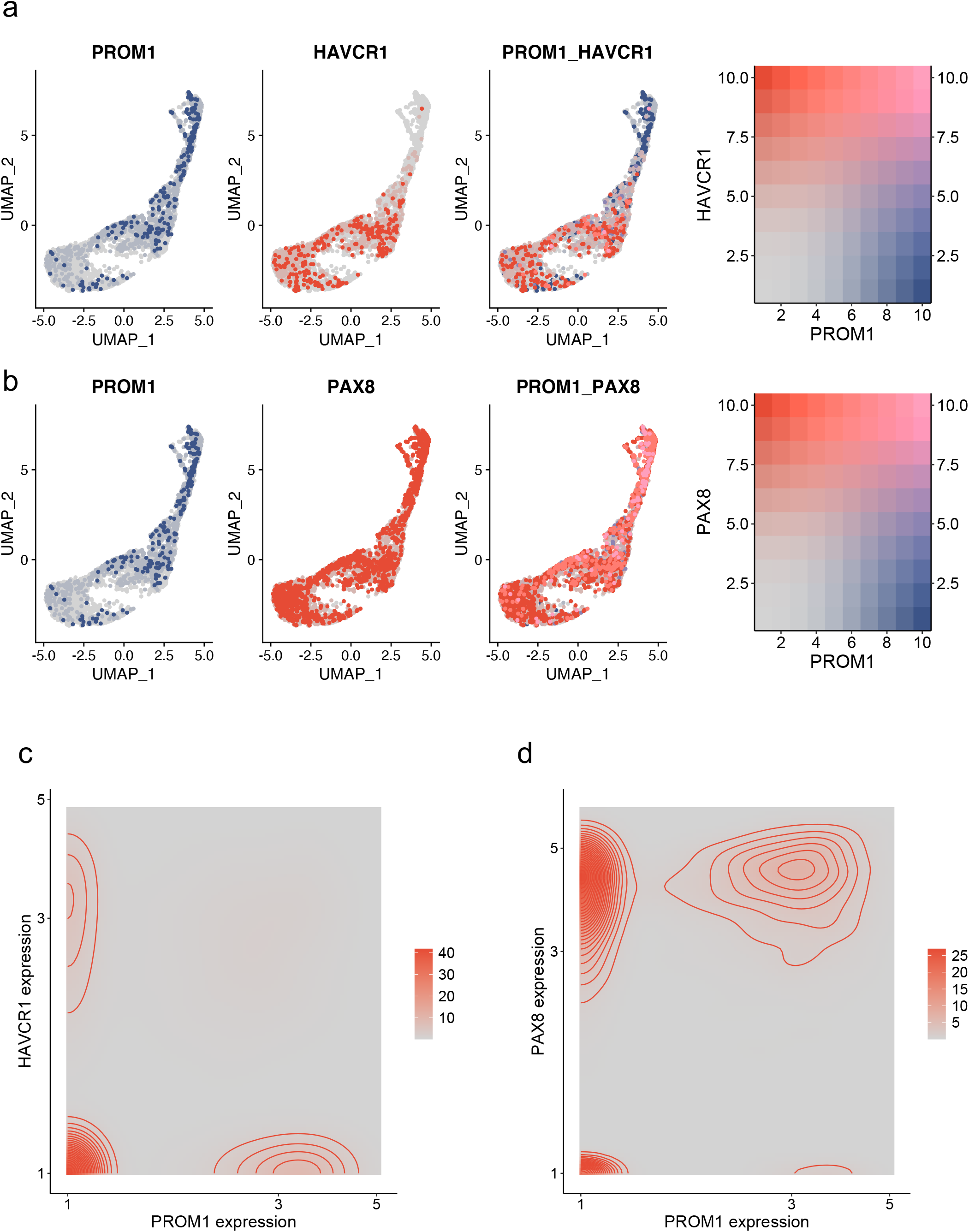
Genes coexpression, a,b) FeaturePlots showing the expression of PROM1, HAVCR1 and coexpression of PROM1-HAVCR1 (a) and PROM1, PAX8 and coexpression of PROM1-PAX8 (b) in the PT cells, c,d) Density plot showing, for each PT cell, the combined expression of PROM1 and HAVCR1 (c) or PAX8 (d), red circles indicating density.

## Methods

### Study approval

All patients admitted to the intensive care unit of the Geneva University Hospitals between April 5^th^ 2020 and May 15^th^ 2020 were screened. Patients were included if they met the following inclusion criteria: positive covid status defined as a positive PCR for SARS-Cov-2 and pneumonia, older than 18, administration of sedative and class III analgesic and if a decision of treatment withdrawal had been taken by the attendings physicians. The consent was given orally by the next of kin, due to the ban on visits in force in our hospital. The study was approved by the local ethical committee for human studies of Geneva, Switzerland (CCER 2020-00644, Commission Cantonale d’Ethique de la Recherche) and performed according to the Declaration of Helsinki principles.

### Kidney biopsies

Once the consent obtained, kidney biopsies were performed by DL and SDS, just before the therapeutic withdrawal. Using 18G Automated biopsy guns (Max-Core, BardCare) in combination with real-time ultrasound guidance, taking five cores’ biopsies were collected from the right kidney. They were directly frozen and stored at −80°C until process.

### Histology and immunochemistry

Human kidney biopsy specimens were fixed with formaldehyde 4%, dehydrated and paraffin-embedded. 2μm kidney sections were stained with hematoxylin and eosin, periodic acid–Schiff, Jones, and Masson’s trichrome for evaluation of tubular injury, inflammatory infiltrate and kidney fibrosis. Immunochemistry staining were performed as followed: after antigen retrieval with pressurized heating chamber in citrate buffer pH7 or tris-EDTA pH9, 5 μm tissue sections were incubated with antibodies mouse monoclonal anti-human HAVCR1 (dilution 1:250, clone 219211, RD Systems), rabbit polyclonal anti-human PROM1 (dilution 1:250, Novus biological) and mouse monoclonal anti-human VCAM1 (dilution 1:25, clone 1.4C3, Invitrogen) for 1h at room temperature. Then, the slides were incubated with the appropriate horseradish peroxidase– conjugated secondary antibody (Dako, Via Real Carpintera, USA) for 30mn at room temperature. Slides were developed using diaminobenzidine chromogen and then counterstained with Mayer hematoxylin. Stained sections were examined with a Zeiss microscope (Zeiss, Oberkochen, Germany). Negative controls were performed in absence of primary antibody.

### Single-nucleus RNA-seq analysis

#### Mouse EDU model

The experimental model was previously characterized (Legouis et al., 2020; Liu et al., 2017). Briefly, 10- to 12-week-old, 25–28-g, male C57BL6/J or Six2TGC Rosa26rtTA pTREH2-GFP mice were anaesthetized with an intraperitoneal injection of ketamine/xylazine (ketamine, 105 mg kg–1; xylazine, 10 mg kg–1). Body temperature was maintained at 36.5–37.0 °C throughout the procedure. The kidneys were exposed by a midline abdominal incision, and both renal pedicles were clamped using non-traumatic micro-aneurysm clips (Roboz Surgical Instrument Co.). Ischemic period was 15 minutes. Restoration of blood flow was monitored by the return of normal kidney colour after removal of the clamps. All mice received 1 ml of normal saline intraperitoneally at the end of the procedure. Sham-operated mice underwent to the same procedure except for clamping of the pedicles. Ethynyldeoxyuridine (0.05 mg g–1; Sigma, Cat#900584) was injected 48 h postoperatively to identify cells proliferating after injury. Kidneys were harvested at 96hr and 28days and snap-frozen with liquid nitrogen.

#### snRNA-seq sample processing

Single nuclei isolation from tissue was performed as previously described (Legouis et al., 2020). Briefly, frozen human sample biopsies or mouse renal tissues were cut into small pieces and transferred into a 2-ml Dounce homogenizer (Sigma, Cat#D8938) loaded with 1 ml of NEZ Lysis Buffer (Sigma, Cat#N3408) with RNase inhibitor (NEB, Cat#M0314) at final concentration of 0.4U/μl on ice. Samples were then Dounce homogenized on ice with five strokes of the looser pestle every 2 min for 8 min (25 strokes in total). Samples were then slowly Dounce homogenized 25 times with the tighter pestle on ice. The homogenized sample was filtered through a 40-μm Falcon Nylon Cell Strainer, then the filter was washed with 8 ml of 1% BSA PBS, and the nuclear suspension spin in a precooled (4 °C) centrifuge at 650g for 8 min. Supernatant was removed, the pellet re-suspended in 2% BSA PBS with RNase inhibitor and nuclei from human biopsies further filtered by 20-μm and 10-μm strainers and moved to a low-bind Eppendorf tube. Nuclei from murine samples were sorted as DAPI and EDU positive on a BD Aria into 2% BSA solution with RNA inhibitor after 40um filtering. Nuclear quality and number were assessed with trypan blue staining.

Single-cell transcriptomes was performed using 10X Chromium single cell platform (10X Genomics) and processed according to the 10X Chromium protocol. Barcoded single-cell gel beads in emulsion (GEMs) were created by 10x Genomics Chromium TM and then reverse transcribed to generate single-cell RNA-seq libraries using Chromium Single Cell 3’ Library and Gel Bead Kit v2 (10X Genomics) according to manufacturer’s instructions. Resulting short fragment libraries were checked for quality and quantity using an Agilent 2100 Bioanalyzer and Invitrogen Qubit Fluorometer. Sequencing Unique molecular identifiers (UMIs), which were incorporated into the 5’ end of cDNA during reverse transcription, were used to quantify the exact number of transcripts in a cell. Paired-end sequencing was carried out on Illumina NextSeq500platform using 150-cycle High Output.

#### snRNA-seq data processing

Sequencing data were processed by CellRanger (version 3.1.0) and reads were aligned to mouse pre-mRNA reference genome (mm10 v3.0.0) and human reference genome (GRCh38-2020). The Cell Ranger *cellranger count* function output filtered gene–cell expression matrices removing cell barcodes not represented in cells. Finally, a UMI count table utilizing both exonic and intronic reads was generated for downstream analysis. The whole data processing was executed by running the script on the Ente Ospedaliero Cantonale server, Switzerland.

Seurat v4 in R v4 was used for downstream analyses, including normalization, scaling, and clustering of nuclei. First, we analyzed each sample separately and excluded nuclei with less than 150 nFeature_RNA detected or more than 4 times the absolute median of nFeature_RNA. We also excluded nuclei with a relatively high percentage of UMIs mapped to mitochondrial genes (>1 and only for one human case >5) and ribosomal genes (>1). Subsequently, we applied SoupX to remove ambient RNA contamination from the human samples. Ambient RNA was estimated from the empty droplet pool with setting “nonExpressedGeneList” to hemoglobin genes (https://github.com/constantAmateur/SoupX). We performed curated doublet removal based on known lineage-specific markers. The samples were integrated to avoid batch effect using Seurat standard work flow split by sample (*orig.ident)* for human samples and split by *dataset* for mouse samples, see **Supplementary Information 1** for a detailed composition (“CovAKI dataset detailed composition.xls”). Following ScaleData, RunPCA, FindNeighbours and FindCluster at a resolution of 0.5 were performed. FindAllMarkers generated the list of genes differentially expressed in each cluster compared to all other cells, within the major subgroups defined (nephron cells, collecting duct, other cells) based on the Wilcoxon rank-sum test and limiting the analysis to upregulated genes with a cut-off for minimum log fold change difference 0.25) and minimum cells with expression (0.1); Results for the whole dataset, the PT subset and the EDU assays are shown in **Supplementary Information 3** (AllSamples_FindAllMarkers.xlsx), **Supplementary Information 4** (PT_Covid_FindAllMarkers.xlsx) and **Supplementary Information 5** (EDU_FindAllMarkers.xls). Cluster reassignment was performed based on manual review of lineage-specific marker expression.

Feature plots were drawn as scatterplots from a given reduction showing the gene-weighted density, using the Nebulosa package.

Secondary Seurat analyses on PT cells employed the SubsetData function to create new r-objects from cohorts of primary analysis. On the subset object, we applied the RunPCA function with default parameters and RunUMAP on 30 dimensions with n.neighbors and min.dist set to 50 and 0.01 respectively. Pseudotime inference was performed iteratively for the three defined trajectories using slingshot (v1.8.0), on the previously clusterized Seurat object. For each trajectory, only the starting cluster was specified.

For each trajectory gene expression among pseudotime was fitted using tradeSeq (v1.5.10), by first running the fitGAM function with 6 knots. The counts matrix, extracted from the Seurat object and the pseudotime inferred by slingshot were used as inputs. The pattern clusterisation was then performed via the clusterExpressionPatterns function, selecting genes that were significantly associated with pseudotime (pvalue <0.05, in the output of the associationTest function).

For each pattern cluster, pathway enrichment was performed using hypeR with a hypergeometric test and the REACTOME, KEGG, GOBP, WIKI, BIOCARTA, and HALLMARK databases downloaded from the Molecular Signature Database (https://www.gsea-msigdb.org/gsea/msigdb/) using the msigdb_gsets function.

The pathway enrichment score of undifferentiated cells was performed with the gsePathway function from the ReactomePA (v1.34.0) package, using a false discovery rate of 0.1 and a minimal geneSet size of 1. The input list of genes was extracted from the Seurat object, with the FindAllMarkers function, using a Wilcoxon test) and sorting according to the log-fold-change variable.

The HeatMap for gene expression among pseudotime was drawn with pheatmap using the scaled fitted genes expression from the clusterExpressionPatterns output.

Score among clusters were calculated as previously described(Macosko et al., 2015) and summarized in the tutorial by P.-Y. Tung (https://jdblischak.github.io/singleCellSeq/analysis/cell-cycle.html), using normalized gene expression from the Seurat object as input and setting the gene correlation value to 0.1. The following genesets were used.

- For the metabolic analyses: GO:0034627, R-HSA-70171, R-HSA-70263, R-HSA-77289, R-HSA-71403, R-HSA-1592230, R-HSA-197264. For gluconeogenesis (R-HSA-70263) and glycolysis (R-HSA-70171) genesets, the shared genes were excluded.
- For the extra cellular matrix, we used the previously described geneset (Kuppe et al., 2021) downloaded from https://raw.githubusercontent.com/mahmoudibrahim/KidneyMap/master/assets/public/ecm_genes_human.txt".

For the proliferation and the pathway activity over time, we followed the same methods but using the scaled fitted genes expression from the clusterExpressionPatterns function as the input matrix and the following list of genes:

- For the S and G2:M scores, the previously described set of genes(Tirosh et al., 2016) for the S and G2M scores.
- For the proliferation index, we used the metaPCNA score previously described (Venet et al., 2011)
- For the NOTCH, HIPPO, CALCIUM and HNF4A pathways, we used the GO:0007219, GO:0035329, GO:0019722 and M11410 genesets, downloaded from the Molecular Signature Database (https://www.gsea-msigdb.org/gsea/msigdb/) using the msigdb_gsets function.

For the WNT pathway activity over time, we used PROGENY (v 1.12.0) with the scaled fitted genes expression from the clusterExpressionPatterns function as the input matrix

For dissimilarity index, we first selected the gene significantly associated with pseudotime for both human and mice datasets (pvalue<0.05 in the AssociationTest output from Tradeseq) and that were shared among species. We then fitted the gene expression along pseudotime with the predictSmooth function from Tradeseq. We finally calculated the Dissimilarity Index, for each pair of pseudotime related fitted gene expression (https://link.springer.com/content/pdf/10.1007/s11634-006-0004-6.pdf) using the TSclust package.

For the heatmap comparing pathways activity in the four clusters, we used PROGENY (v 1.12.0) or the score described above with gene sets from the REACTOME database or the proliferation index and the normalized gene expression from the Seurat object as input. The mean by cluster was thus calculated and shown in a heatmap using pheatmap and row scaling. We used also PROGENY (v 1.12.0) for the allograft kidney recipients RNAseq database. The raw counts were first normalized with the TMM method from the EdgeR package and expressed as count per million. The resulting matrix was used as an input in Progeny. The comparisons between pre and post transplantation samples were performed with a Wilcoxon test.

#### Public snRNA-Seq dataset integrated

GEO datasets were downloaded from Gene Expression Omnibus (http://www.ncbi.nlm.nih.gov/geo/). GSE131882 comprises 3 early human diabetic kidney samples and 3 controls characterized by single nucleus RNA sequencing(Wilson et al., 2019). Control samples represent non-tumor tissue in patients undergoing nephrectomy for renal mass. The control samples were integrated to the COVID-19 dataset. GSE139107 includes a sum of 24 renal samples characterized by snRNA-seq as previously published (Kirita et al., 2020). Briefly, bilateral IRI was induced for 18 minutes in C57B6/J 8- to 10-wk-old male mice and body temperature was monitored and maintained at 36.5 to 37.5 °C throughout the procedure. Kidney were harvested at 0,4,12,48 hrs, 14 days and 6 weeks post injury and analyzed by snRNA-seq using 10X Genomics. The time points: 0,4,12 hrs have been selected and used to validated the dedifferentiation trajectory.

GSE151167 includes 5 samples characterized by snRNA-seq as previously described (Legouis et al., 2020) and reported in this paper. GSE151167 dataset was integrated to the EDU dataset generated in this paper.

## Data and Code Availability

The new snRNA-seq data generated in this study will be deposited on GEO and the accession numbers will be included here. The public datasets used are from GEO with the accession numbers: GSE131882 (Wilson et al., 2019), GSE139107(Kirita et al., 2020), GSE151167(Legouis et al., 2020).

